# Temporal evolution and adaptation of SARS-COV-2 codon usage

**DOI:** 10.1101/2020.05.29.123976

**Authors:** Elisa Posani, Maddalena Dilucca, Sergio Forcelloni, Athanasia Pavlopoulou, Alexandros G. Georgakilas, Andrea Giansanti

## Abstract

The outbreak of severe acute respiratory syndrome-coronavirus-2 (SARS-CoV-2) has caused an unprecedented pandemic. Since the first sequenced whole-genome of SARS-CoV-2 on January 2020, the identification of its genetic variants has become crucial in tracking and evaluating their spread across the globe.

In this study, we compared 134,905 SARS-CoV-2 genomes isolated from all affected countries since the outbreak of this novel coronavirus with the first sequenced genome in Wuhan, China to quantify the evolutionary divergence of SARS-CoV-2. Thus, we compared the codon usage patterns of SARS-CoV-2 genes encoding the membrane protein (M), envelope (E), spike surface glycoprotein (S), nucleoprotein (N), RNA-dependent RNA polymerase (RdRp). The polyproteins ORF1a and ORF1b were examined separately.

We found that SARS-CoV-2 tends to diverge over time by accumulating mutations on its genome and, specifically, on the sequences encoding proteins N and S. Interestingly, different patterns of codon usage were observed among these genes. Genes *S* and *N* tend to use a narrower set of synonymous codons that are better optimized to the human host. Conversely, genes *E* and *M* consistently use the broader set of synonymous codons, which does not vary in respect to the reference genome. CAI and SiD time evolutions show a tendency to decrease that emerge for most genes. Forsdyke plots are used to study the nature of mutations and they show a rapid evolutionary divergence of each gene, due to the low values of x-intercepets.

## 1 Introduction

The recent emergence of the novel, human pathogen Severe Acute Respiratory Syndrome Coronavirus 2 (SARS-CoV-2) in China and its rapid spread poses a global health emergency. On March 11 2020, WHO publicly declared the SARS-CoV-2 outbreak as a pandemic. SARS-CoV2 belongs to *Coronaviridae* family, with a genome around 30 kb in length and a structure characteristic of known coronaviruses. The virus genome encodes for structural, non-structural, and accessory proteins [12]. The four structural proteins are the envelope protein (E), the membrane protein (M), the nucleocapsid protein (N), and the spike glycoprotein (S). ORF1ab gene encodes for the two isoforms of the poliprotein complex pp1a and pp1ab, where the first is the intial segment of the latter, which are further proteolytically cleaved to produce 16 different non-structural proteins. [1], [2], [3].

The nucleocapsid protein plays an important role in maintaining the RNA conformation stable for the replication, transcription, and translation of the viral genome along with protecting the viral genome [9]. It is highly immunogenic and capable of modulating the metabolism of an infected cell [9]. The envelope protein acts as a viroporin [15] and plays multiple roles in viral replication and signaling pathways that affect inflammatory and type 1 INF *γ* signaling [13]. The spike protein *S* is responsible for receptor recognition and membrane fusion that leads to viral entry into the host cells [23]. Finally, the membrane protein is associated with the spike protein and is responsible for the virus budding process [8].

Characterization of viral mutations can provide valuable information for assessing the mechanisms linked to pathogenesis, immune evasion and viral drug resistance. In addition, viral mutation studies can be crucial for the design of new vaccines, antiviral drugs and diagnostic tests. The mutation rate of RNA viruses (10^*−*5^*−*10^*−*3^ per replication cycle [10]) is dramatically higher than their mammalian hosts (10^*−*9^ per year [11]). This high mutation rate is correlated with virulence modulation and evolvability, which are considered beneficial for viral adaptation [4].

According to data from the public database Global Initiative on Sharing All Influenza Data (GISAID), seven major clades of SARS-CoV-2 can be identified and are referred to as clade V (variant of the ORF3a-G251V and of NSP6-L37F), and clade S (variant ORF8-L84S and NSP4:S76S), clade G (variant of the spike protein S-D614G and of NSP12b-P314L), and the subclades GH and GR, that have an additional mutation to the G ones (respectively, ORF3a-Q57H and N-RG203KR), and finally the GV that presents both G and V mutations [6, 7]. In particular, Giorgi et al., showed that clade G, the most spread of all clades but prevalent in Europe, carries a D614G mutation in the Spike protein, which is responsible for the initial interaction of the virus with the host human cells [7].

In the present study, we investigated the evolution of SARS-CoV-2 genomes and codon usage patterns over time, as well as virus adaptation to the human host. For this purpose, we focused on the four SARS-CoV-2 genes encoding the structural proteins membrane (M), envelope (E), spike surface glycoprotein (S) and nucleoprotein (N), as well as RNA-dependent RNA polymerase (RdRp) and the two segments of polyproteins ORF1ab separately, ORF1a and ORF1b.

## 2 Materials and Methods

### 2.1 Sequence data analyzed

All available SARS-CoV-2 genomes reported across the world were obtained from GISAID (available at https://www.gisaid.org/epiflu-applications/nexthcov-19-app/), on December 4, 2020. Then, the sequences were classified according to their isolation dates. Only complete genomes (29-30 Kb) were included in the present analysis, with a list of 134,905 SARS-CoV-2 genomes. We used the SARS-CoV-2 coding DNA sequences (CDSs) deposited in January 2020 by Zhu and coworkers [14], formerly called “Wuhan seafood market pneumonia virus”, also referred to as “Wuhan-Hu-1” (WSM, NC 045512.2), as reference sequence. We retrieved these sequences from NCBI public database at https://www.ncbi.nlm.nih.gov/. The coding DNA sequences (CDS) of the reference SARS CoV-2 genome (NC 045512.2) were used to retrieve the homologous protein-coding sequences from the genomes under study, by using Exonerate with default parameters [16].

### 2.2 Relative Synonymous Codon Usage

*RSCU* vectors for all the genomes were computed by using an in-house Python script, following the formula:

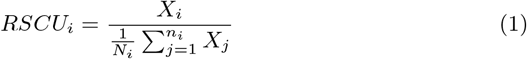

In the *RSCU*_*i*_ *X*_*i*_ is the number of occurrences, in a given genome, of codon i, and the sum in the denominator runs over its *n*_*i*_ synonymous codons. If the *RSCU* value for a codon *i* is equal to 1, this codon was chosen equally and randomly. Codons with *RSCU* values greater than 1 have positive codon usage bias, while those with value less than 1 have relatively negative codon usage bias [22].

Stop codons and non-degenerate amino acids have been excluded from the calculations, making RSCU a 59-component vector.

### 2.3 Principal Component Analysis

Principal Component Analysis (PCA) [38] is a multivariate statistical method to transform a set of observations of possibly correlated variables into a set of linearly uncorrelated variables (called principal components) spanning a space of lower dimensionality. The transformation is defined so that the first principal component accounts for the largest possible variance of the data, and each subsequent component in turn has the highest variance possible under the constraint that it is orthogonal to (*i*.*e*., uncorrelated with) the preceding components.

We use this technique on the space of *RSCU* vectors, so that each gene of SARS-COV-2 is represented as a 59-dimensional vector with codons as coordinates. Such coordinates are separately normalized to zero mean and unit variance over the whole genome. We then obtain the associated covariance matrix between the four dimensions of codon bias and diagonalize it. The eigenvectors of the covariance matrix, ordered according to the magnitude of the corresponding eigenvalues, are the principal components of the original data.

PCA has been performed by using Pythons sklearn.

### 2.4 Codon Adaptation Index

The codon adaptation index (*CAI*) [21, 22] was used to quantify the extent of codon usage adaptation of SARS-CoV-2 to the human coding sequences. The principle behind *CAI* is that the codon usage in highly expressed genes can reveal the optimal (i.e., most efficient for translation) codons for each amino acid. Hence, *CAI* is calculated based on a reference set of highly expressed genes to assess, for each codon *i*, the relative synonymous codon usages (*RSCU*_*i*_) and the relative codon adaptiveness (*w*_*i*_):

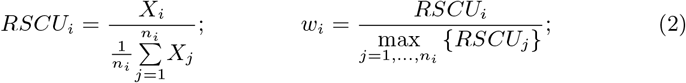

In the *RSCU*_*i*_, *X*_*i*_ is the number of occurrences of codon *i* in the genome, and the sum in the denominator runs over the *n*_*i*_ synonymous codons. *RSCU* measures codon usage bias within a family of synonymous codons. *w*_*i*_ is defined as the usage frequency of codon *i* compared to that of the optimal codon for the same amino acid encoded by *i* (i.e., the the most used one in a reference set of highly expressed genes). Finally, the *CAI* value for a given gene *g* is calculated as the geometric mean of the usage frequencies of codons in that gene, normalized to the maximum *CAI* value possible for a gene with the same amino acid composition:

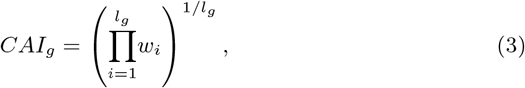

where the product runs over the *l*_*g*_ codons belonging to that gene (except the stop codon).

This index ranges from 0 to 1, where the score 1 represents a greater tendency of the gene to use optimal codons in the host organism. The CAI analysis was performed using *DAMBE* 5.0 [20].

### 2.5 Similarity Index

The similarity index (SiD) was used to provide a measure of similarity in codon usage between SARS-CoV-2 and various potential host genomes. Formally, SiD is defined as follows:

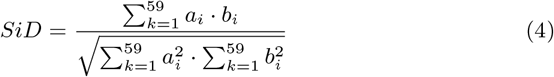

where *a*_*i*_ is the RSCU value of 59 synonymous codons of the SARS-CoV-2 coding sequences; *b*_*i*_ is the RSCU value of the identical codons of the potential host. SiD is defined as the cosine value of the angle included between A and B spatial vectors, and therefore, quantifies the degree of similarity between the virus and the host in terms of their codon usage patterns. In our analysis, we considered https://cubap.byu.edu/) [18]. SiD values range from 0 to 1; the higher the value of SiD, the more adapted the codon usage of SARS-CoV-2 to the host [25].

### 2.6 Effective Number of Codons

We calculated the effective number of codons (*ENC*) to estimate the extent of the codon usage bias of SARS-CoV-2 genes. The values of *ENC* range from 23 (when just one codon is used for each amino acid) to 61 (when all synonymous codons are equally used for each amino acid) [19]. For each sequence, the computation of *ENC* starts from *F*_*α*_, a quantity defined for each family *α* of synonymous codons:

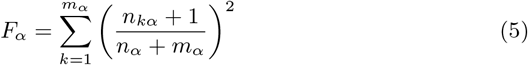

where *m*_*α*_ is the number of different codons in *α* (each one appearing 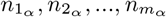 times in the sequence) and 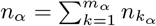

Finally, the gene-specific ENC is defined as:

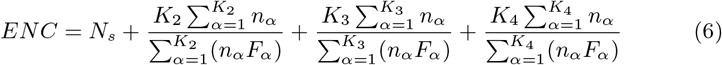

where *N*_*S*_ is the number of families with one codon only and *K*_*m*_ is the number of families with degeneracy *m* (the set of 6 synonymous codons for *Leu* can be split into one family with degeneracy 2, similar to that of phenylalanine (Phe), and one family with degeneracy 4, similar to that, for example, of proline (Pro)). The same applies to the other 6-fold families.

*ENC* was estimated by using a Python script following [17].

### 2.7 ENC plot

An *ENC* plot analysis was performed to estimate the relative contributions of mutational bias and natural selection in shaping CUB of 13 genes encoding proteins that are crucial for SARS-CoV-2. In this plot, the *ENC* values are plotted against *GC*_3_ values. If codon usage is dominated by the mutational bias, then a clear relationship is expected between *ENC* and *GC*_3_:

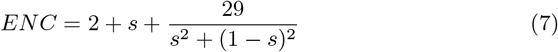

*s* represents the value of *GC*_3_ [19]. If the mutational bias is the main force affecting CUB of the genes, the corresponding points will fall near Wright’s theoretical curve. Conversely, if CUB is mainly affected by natural selection, the corresponding points will fall considerably below Wright’s theoretical curve.

To quantify the relative extent of the natural selection, for each gene, we calculated the Euclidean distance *d* of its point from the curve.

We then showed the average values of the distance over time with a heatmap, drawn with MATLAB. We consider the difference of the distance of the dots from the curve and that of Wuhan-Hu-1 reference genes.

### 2.8 Forsdyke plot

To study the mutational rates of genes, we performed an analysis by using our previously defined Forsdyke plot [43]. Each gene of the SARS-CoV-2 sequencings considered was compared to its orthologous gene in the reference genes of Wuhan-Hu-1 genome. Each pair of orthologous genes is represented by a point in the Forsdyke plot, where protein divergence is correlated with RNA divergence (see Methods in [43] for details). The protein sequences were aligned using Biopython. The RNA sequences were then aligned using the protein alignments as templates.

Then, both RNA and protein divergences were assessed as explained in Methods in [43] by counting the number of mismatches in each pair of aligned sequences. Thus, each point in the Forsdyke plot measures the divergence between pairs of orthologous genes in the two viruses, as projected along with the phenotypic (protein) and nucleotidic (RNA) axis. The first step in each comparison is to compute the regression line between protein vs RNA sequence divergence in the Forsdyke plot getting values of intercept and slope for each variant of genes. To test whether the regression parameters associated with each variant are different or not, we have followed a protocol founded by Dilucca et al., considering a p-value *≤* 0.05.

### 2.9 Protein-Protein Network Analysis

In this study, we used the 332 high-confidence SARS-CoV-2-human protein-protein interactions (PPI) collected by Gordon et al. [31] who identified the viral proteins that physically associate with human proteins using affinity-purification mass spectrometry (AP-MS). We downloaded these PPI from NDEx (https://public.ndexbio.org/network/43803262 *−* 6*d*69 *−* 11*ea − bfdc −* 0*ac*135*e*8*bacf*).

To detect communities of PPI, we used the application Molecular Complex Detection (MCODE) [32] in Cytoscape (https://cytoscape.org/). In a nutshell, MCODE iteratively groups together neighboring nodes with similar values of the core-clustering coefficient, which for each node is defined as the density of the highest *k*-core of its immediate neighborhood times *k*.^1^ MCODE detects the densest regions of the network and assigns to each detected community a score that is its internal link density times the number of nodes belonging to it.

We also characterized the first ten communities *c* with the mean value 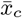 and standard deviation *σ*_*c*_ of codon bias values within the community, and use them to compute a *Z*-score as 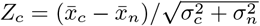 (where 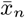 and *σ*_*n*_ are, respectively, the mean value and standard deviation of codon bias values computed for all proteins). In this way, a value of *Z*_*c*_ *>* 1 (*Z*_*c*_ *< −*1) indicates that community *c* features significantly higher (lower) codon bias than the population mean. Cytoscape was used to detect the degree *k* of a protein.

## 3 Results

### 3.1 Dataset Distribution

The records of virus genomes according to geographical location and month of isolation are shown in Figure 1. A detailed report of all extracted genes per region can be found in Table 1.

**Table 1.** Summary of total sequences available per protein, subdivided in geographical regions.

| Continent | Total seqs | ORF1a | ORF1b | RdRp | S | N | M | E |
| --- | --- | --- | --- | --- | --- | --- | --- | --- |
| Africa | 1 914 | 1 796 | 1 689 | 1 904 | 1 780 | 1 878 | 1 914 | 1 913 |
| Asia | 8 148 | 7 764 | 7 752 | 8 117 | 7 975 | 8 069 | 8 148 | 8 127 |
| Europe | 83 108 | 75 772 | 75 549 | 82 895 | 76 156 | 82 784 | 83 101 | 83 107 |
| North America | 30 227 | 28 265 | 27 715 | 30 007 | 27 899 | 30 082 | 30 187 | 30 224 |
| South America | 1 350 | 1 275 | 1 263 | 1 342 | 1 306 | 1 344 | 1 350 | 1 350 |
| Oceania | 10 064 | 8 925 | 9 987 | 10 057 | 8 554 | 9 954 | 10 062 | 10 064 |
| **Total** | **134 811** | **123 797** | **123 955** | **134 322** | **123 670** | **134 111** | **134 762** | **134 785** |

**Figure 1.**
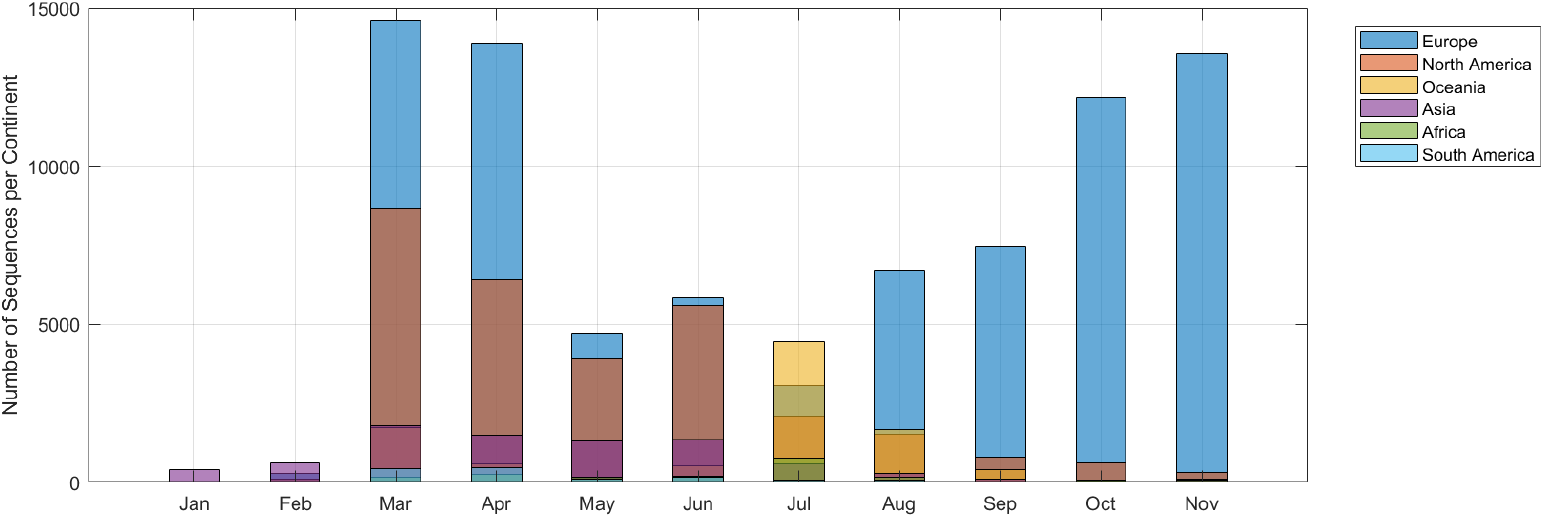
Distribution of the 134,905 genomes used in this study. Distribution of raw sequences per month and geographical region. Details for each gene are summarized in Table 1.

A great percentage of the annotated SARS-CoV-2 genomes (about 62% and 22% respectively) are distributed in Europe and North America. The number of complete viral genomes is non-homogeneous, going from 200-300 in the first months, to a peak of 24,000 in March. In the other months the values fluctuate among 7,000 and 23,000.

### 3.2 Principal Component Analysis of RSCUs

We performed PCA over the space of *RSCU* vectors measured for each SARS-COV-2 gene in all sequences. The two first principal components (*PC*_1_ and *PC*_2_) appear to represent as much as 78% of the total variance of codon bias over the genome (respectively 52% for *PC*_1_ and 26% for *PC*_2_).

The projections of the first two principal components on the individual codons show that none of the codons predominantly contributes to the data variability (see Figure 4). Thus, the placement of a gene on the *PC*_1_-*PC*_2_ plane depends on a weighted contribution of all codons.

A great number of codons highly affect *PC*_1_, resulting in distinct clusters, whereas *PC*_2_ is mostly affected by triplets that code for Phenylalanine (TTT and TTC), Leucine (TTA, TTG and CTA), Isoleucine (ATT, ATC), Lysine (AAG) and Valine (GTA).

Interestingly, the genes encoding for the polyproteins and spike protein are very close in PCA plane in the bottom of the graph, far apart from the others. Regarding *N*, the coordinate in *PC*_1_ is similar to those of the forementioned nucleus, while is quite distant in *PC*_2_. *M* is closer to *N* in *PC*_2_, denoting a similar usage of codons coding for the five amino acids displayed above. Finally, *E* is the most separated cluster due to its shorteness, and apparently affect the RSCU values (see Figure 2).

**Figure 2.**
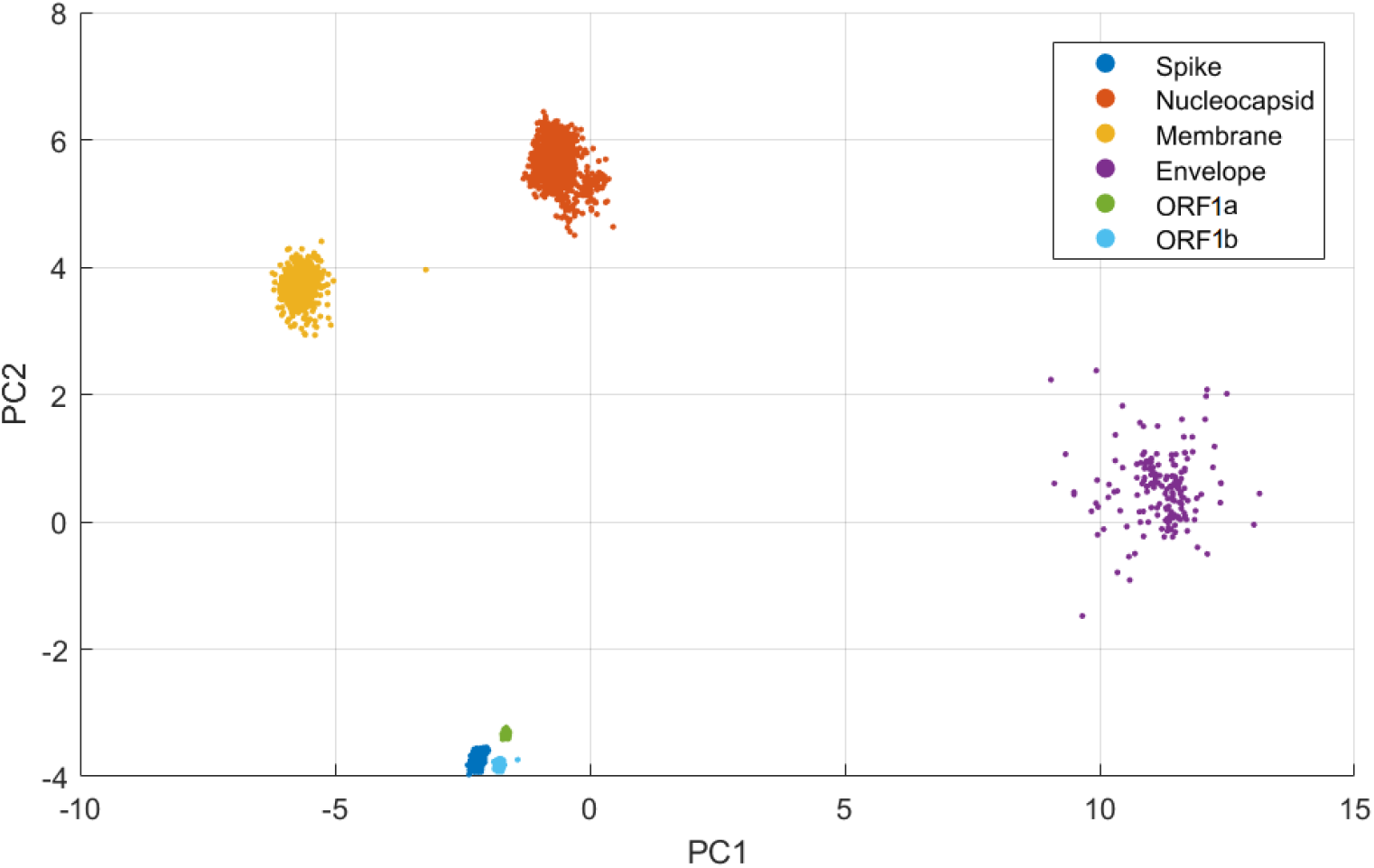
PCA of SARS-CoV-2 Genes. All available sequences for each gene has been used in this study, by calculating RSCUs and performing PCA for each of them. Approximately the same number of data was analyzed for each gene.

**Figure 3.**
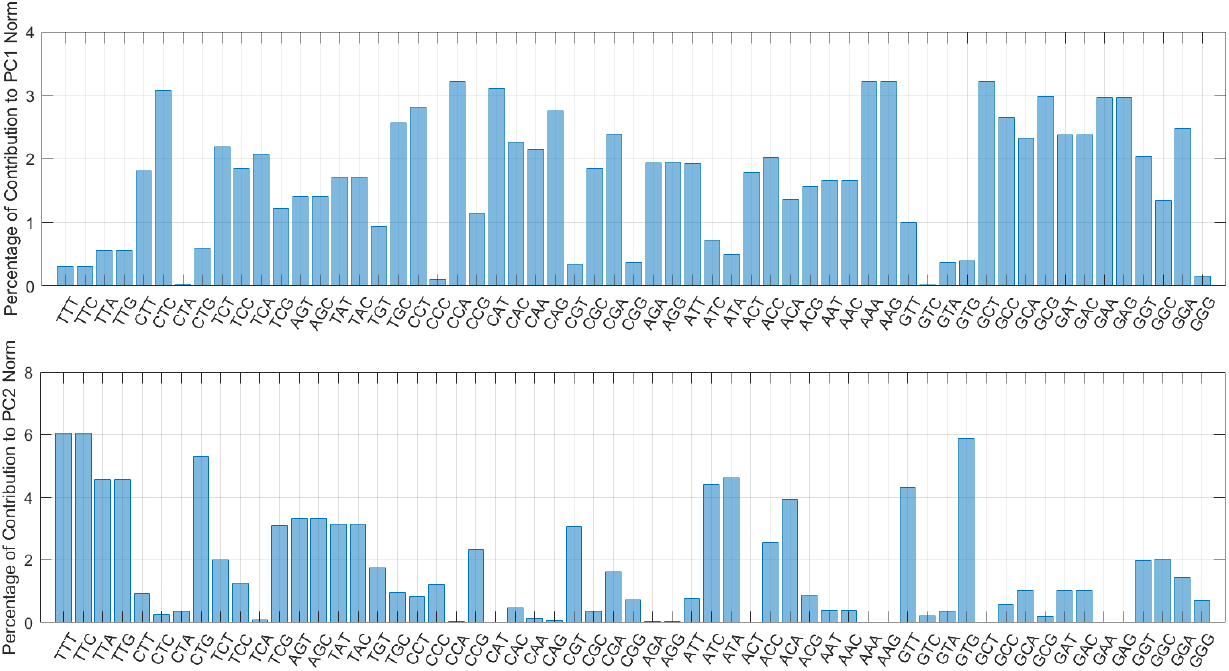
Projection of the first two PCA components on the RSCU vectors. The distribution of each codon in *PC*_1_ is more uniform, resulting from a weighted and coherent contribution of all codons, whereas, in *PC*_2_ the contribution is more focused on specific codons.

**Figure 4.**
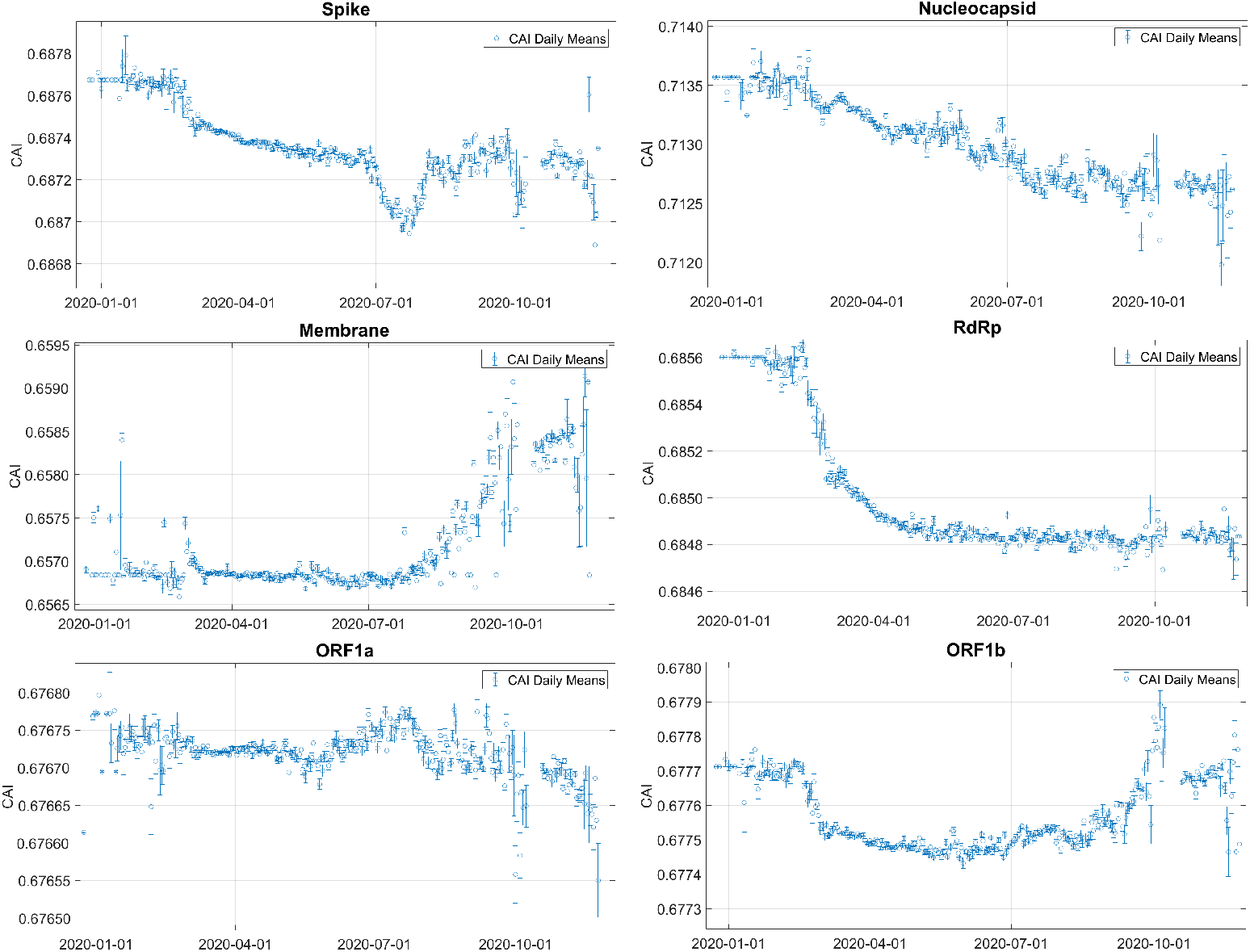
CAI values daily means in SARS-CoV-2 genes. CAI values estimated from all the available sequences, averaged over each day, with standard deviation as errorbar.

Taken together, this analysis shows that the longer genes carry a signature RSCU of the virus, that in two proteins, namely *M* and *N*, differs in the codons encoding for 5 amino acids.

### 3.3 Codon Adaptation Index Time Evolution

To measure the codon usage bias in the SARS-CoV-2 genomes, we first evaluated CAI for all genes under investigation. For each gene, we performed a daily mean of the values, which are displayed in Figure 4. As in the previous analysis, the CAI starting values are very similar for Spike, ORF1a, ORF1b and RdRp while Nucleocapsid has the highest and Membrane shows the lowest value. Envelope protein analysis is redundant for constant CAI values through evolution.

The fastest evolving gene appears to be the one coding for Nucleocapsid with a globally descending trend. Spike too displays the same trend but with a down-peak in the month of July. This is mostly due to Oceania sequences which were very abundant in the summer period. A detailed output of CAI temporal evolution per country can be found for this protein in Figure 6.

Membrane shows a constant evolution until August, where a steep ascent occurs. On the other hand, RdRp has a sudden decrease in the first months and a subsequent stabilization on a lower value. This may be due to the initial adaptation to the new host and the finding of a more optimized amino acidic sequence, that rapidly superseded the older one.

Finally, the two segments of gene ORF1ab manifest a more localized evolution, in accordance with the fundamental role of these proteins for replication.

Overall, the genes display a descending trend, against the hypothesis that a virus evolves in order to align its codon usage bias to the one of the host. This phenomenon though, has previously been observed by Dilucca et al. [24] and is thought to be beneficial for the virus, as it can exploit the host resources at its best. A plausible explanation is that the virus has already adapted its genome to *Homo Sapiens*. In order to spread, a virus “is willing” to keep the host alive, and therefore they adapt a sub-optimal codon usage bias. If the virus replicates efficiently it is more likely to be more successful at invading a given host, and, eventually kill it.

### 3.4 SiD

In line with our previous study [24], we calculated the similarity index (SiD) of SARS-CoV-2 genomes in respect to the human host. To understand the rationale behind these results, the higher the value of SiD, the more adapted the codon usage of SARS-CoV-2 to the host under study [25].

In the present analysis, we calculated SiD for each gene under study, for all the available sequences, and performed a daily average (see Figure 5).

**Figure 5.**
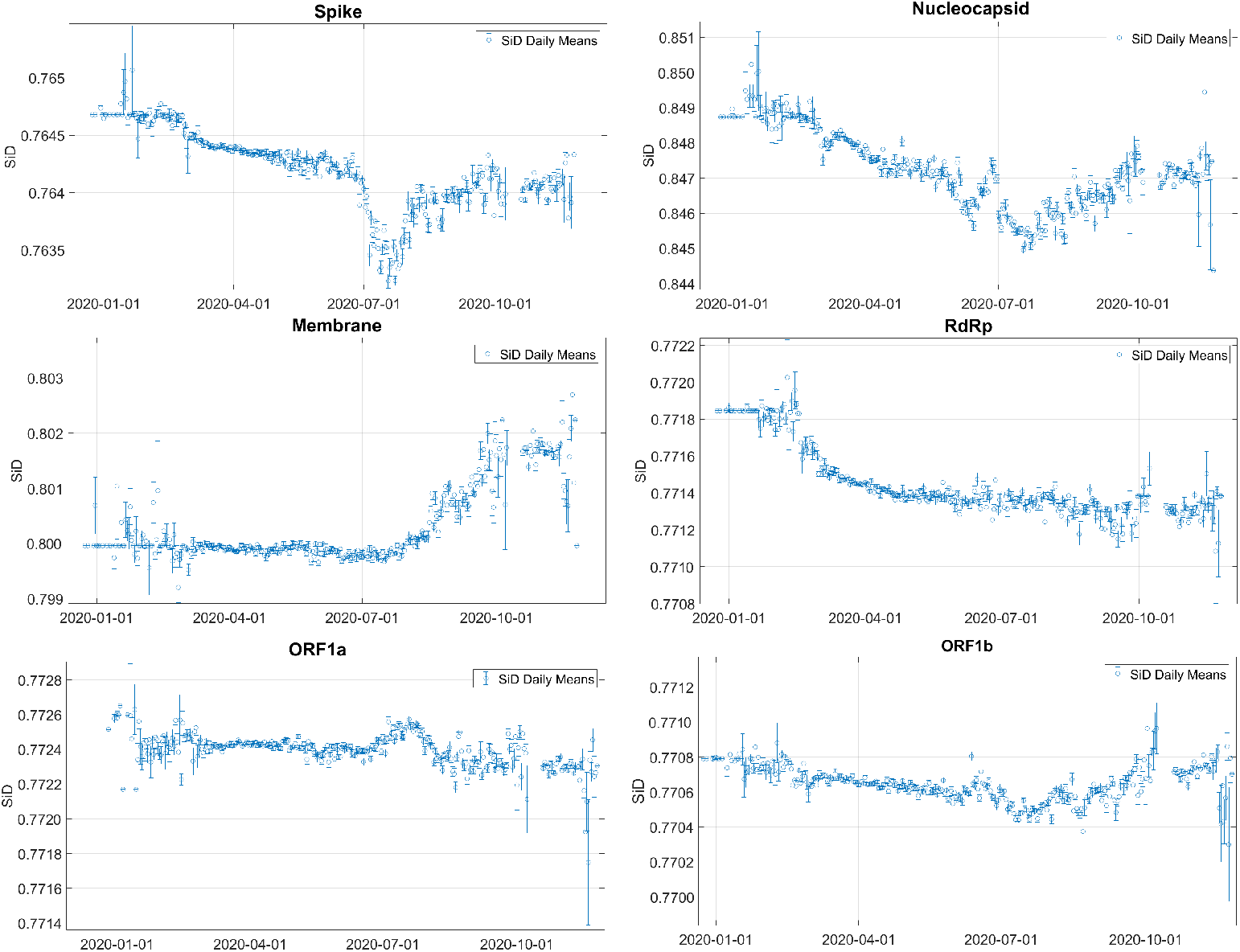
SiD values daily means in SARS-CoV-2 genes. SiD values estimated from all the available sequences, averaged over each day, with standard deviation as error bar.

**Figure 6.**
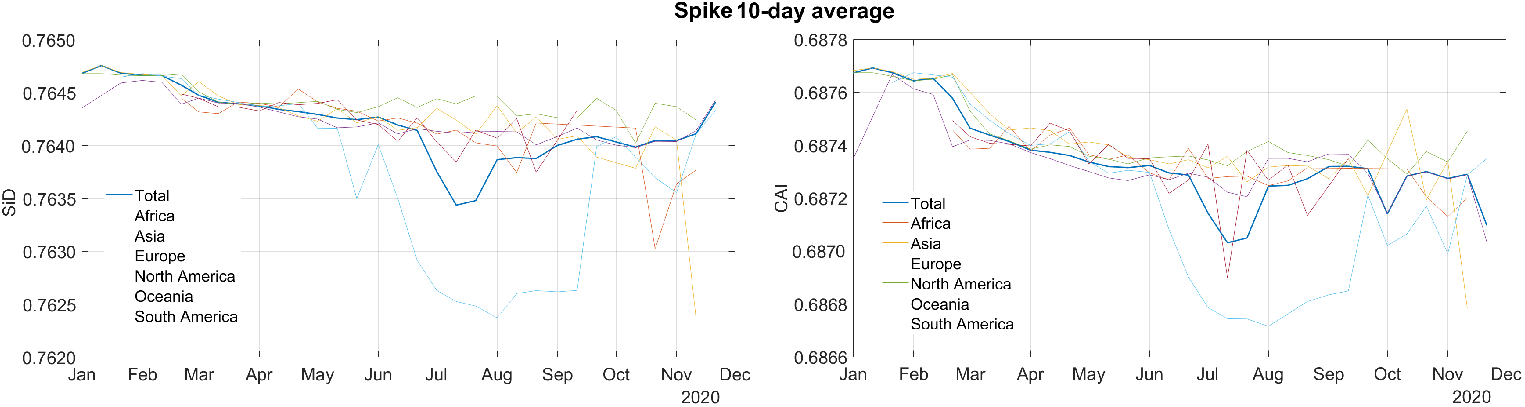
SiD and CAI values region details. SiD and CAI temporal evolution per country can be found for Spike protein with an average every 10 days to explain the down-peak occuring in July.

In line with the previous statements concerning CAI, the more adapted gene appears to be the one coding for Nucleocapsid, while the gene encoding Spike seems to be the least adapted. Also in this case Spike, RdRp and ORF1ab show similar values, confirming the hypothesis of a signature codon bias.

Moreover, the trends are mostly descending, remarking the previous statements on CAI.

By looking at the Spike temporal evolution of SiD, it shows the down peak in July, as occurred for CAI. To have a clearer picture we made a region-based analysis, averaging over 10 days to smooth the curves. It is immediate from Figure 6 that the peaks for both CAI and SiD are due to sequences from Oceania.

### 3.5 ENC Plot Analysis of SARS-CoV-2 genes

To further investigate the evolutionary forces that affect the SARS-CoV-2 codon usage, an ENC-plot analysis was conducted separately for each of the genes considered herein. First, a scatter plot of ENC vs *GC*_3_ of reference sequence Wuhan-Hu-1 genes has been created, together with the curve, in Figure 7. This was done to have a reference for the subsequent temporal evolution. All genes are found below the curve, meaning that they have evolved under a selective pressure [19].

**Figure 7.**
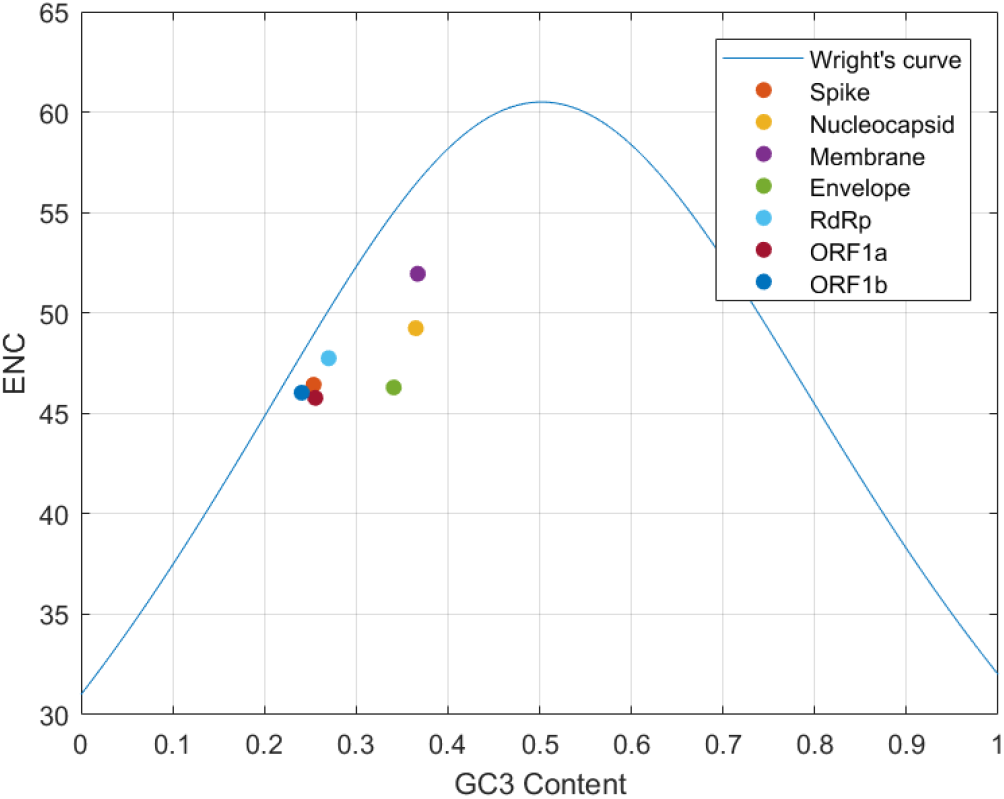
ENC plot of Wuhan-Hu-1 genes. Representation of Wright’s Theoretical curve together with Wuhan-Hu-1 genes ENC Plots (*GC*_3_ vs *ENC*).

Afterwards variations over time have been investigated by generating ENC plots for sets of genes binned by the time. To illustrate more clearly the temporal diversities, 10-day averages of the distances from Wright’s theoretical curve have been estimated. Then we calculated the difference between these values and the scattered values of Wuhan-Hu-1, represented in Figure 7 and graphically illustrated in a heatmap in Figure 8.

**Figure 8.**
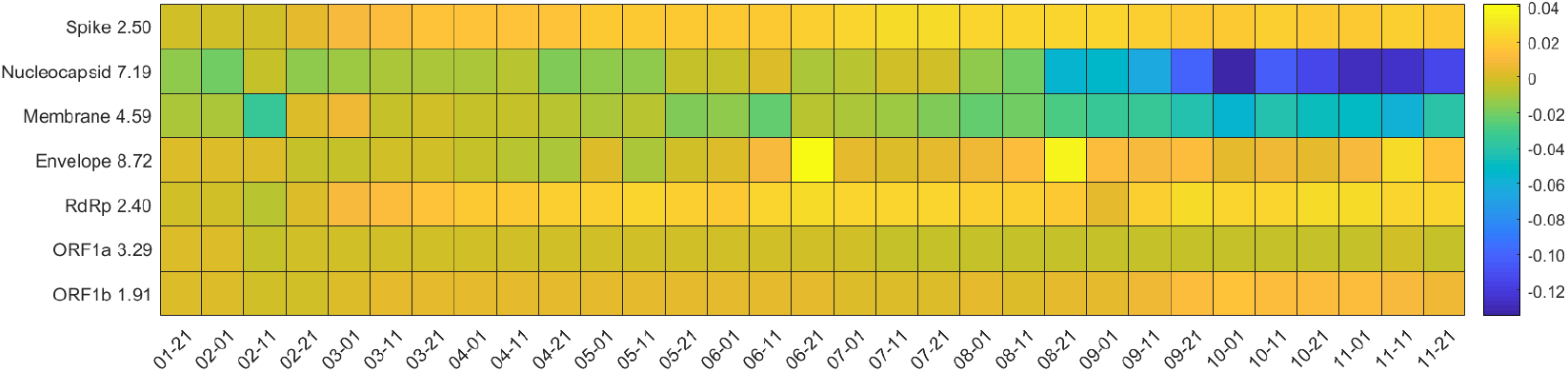
Heatmap of ENC Plots. Each square of the heatmap represents the distance between the reference gene distance from Wright’s Curve and the examined gene one. 10-day averages have been calculated. All dates refer to year 2020.

The average distances from Wright theoretical curve tend to increase over time for most genes, except for ORF1a, which appears to be stable. This could be interpreted as the action of natural selection over time. On the other hand, Nucleocapsid and Membrane are getting closer to the curve, suggesting that the codon usage of these genes is mostly affected by mutational bias. It should be mentioned though that, except for E that has a different bias due to its short length, M and N have the greatest distance from Wright’s curve, pointing that they already are the most subject to natural selection.

### 3.6 Forsdyke plots

The RNA and protein sequence divergence of the seven selected genes have been analyzed by comparing the nucleotide sequences of the reference Wuhan-Hu-1 genes and their corresponding protein sequences with the other SARS-CoV-2 sequences under study. This has been done to estimate evolutionary divergences of each gene. Each gene is represented in a different panel in Figure 9. Usually each sequence is represented by a point in the plot but due to the discreetness of mutations and redundancy, the points were often superimposed. To give an idea of the number of data with the same coordinates, a bubble plot has been made, with the radius of each circle proportional to the degeneracy of the point.

**Figure 9.**
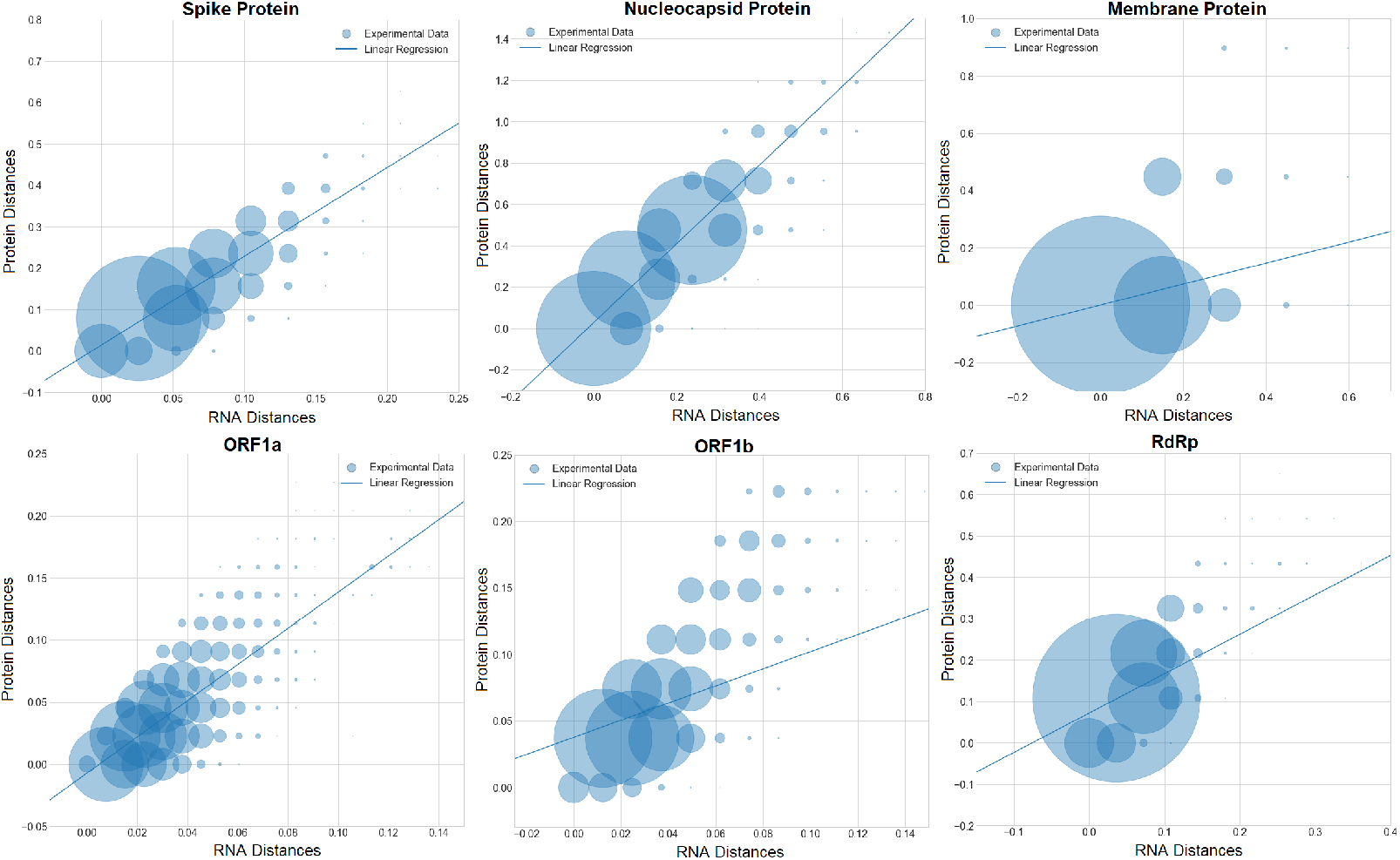
Forsdyke plots of the considered genes. Forsdyke Plots of structural proteins (upper row) and replicase polyprotein complex (lower row). The radius of each circle is proportional to the number of data points in the same spot. The inferred data from linear regression can be found in Table 2.

Each point in the Forsdyke plots represents the divergence between a pair of orthologous genes - in this case being part of the same species, they are variants of the same gene - as projected along the phenotypic (protein) and nucleotide (RNA) axis. Thus, the slope is an estimation of the fraction of RNA mutations that results in amino acid substitutions [43]. In Figure 9, a separate Forsdyke plot is shown for each gene. Envelope protein plot did not show many mutations.

**Table 2.**
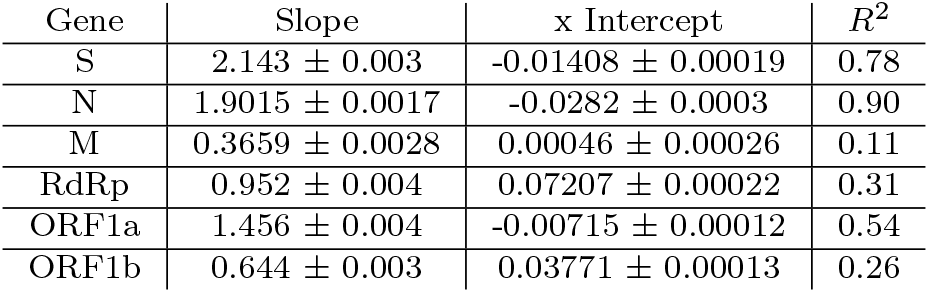
Forsdyke Plots Linear Regressions. Linear regressions parameters of Forsdyke plots in Figure 9.

| Gene | Slope | x Intercept | $R^2$ |
| --- | --- | --- | --- |
| S | $2.143 \pm 0.003$ | $-0.01408 \pm 0.00019$ | 0.78 |
| N | $1.9015 \pm 0.0017$ | $-0.0282 \pm 0.0003$ | 0.90 |
| M | $0.3659 \pm 0.0028$ | $0.00046 \pm 0.00026$ | 0.11 |
| RdRp | $0.952 \pm 0.004$ | $0.07207 \pm 0.00022$ | 0.31 |
| ORF1a | $1.456 \pm 0.004$ | $-0.00715 \pm 0.00012$ | 0.54 |
| ORF1b | $0.644 \pm 0.003$ | $0.03771 \pm 0.00013$ | 0.26 |

**Table 3.** Features of ten top-scoring communities. Number of nodes (*n*), community score (*n* times the internal density), mean degree, average codon bias indices (CAI and ENC).

| ID | $n$ | score | k | CAI | ENC |
| --- | --- | --- | --- | --- | --- |
| 1 | 13 | 13.00 | 14.15 | 0.77 | 51.99 |
| 2 | 6 | 6.00 | 5.67 | 0.69 | 55.88 |
| 3 | 6 | 6.00 | 6.83 | 0.71 | 53.24 |
| 4 | 5 | 5.00 | 4.80 | 0.70 | 53.70 |
| 5 | 5 | 5.00 | 4.40 | 0.70 | 53.05 |
| 6 | 4 | 4.00 | 4.00 | 0.71 | 53.11 |
| 7 | 4 | 4.00 | 4.00 | 0.70 | 53.12 |
| 8 | 4 | 4.00 | 3.25 | 0.72 | 53.12 |
| 9 | 4 | 4.00 | 6.25 | 0.70 | 53.10 |
| 10 | 4 | 3.33 | 3.50 | 0.72 | 53.13 |

Overall, protein and RNA sequence divergences are linearly correlated, and these correlations correspond to slopes and intercepts of the linear regressions in Table 2.

Gene M displays the lowest slope, pointing that this protein tends to evolve slowly by accumulating mutations on its corresponding gene. The low value of the R-squared is due to the fact that it is the shortest protein and it has less mutations, thereby appearing more scattered in the plot. Conversely, the steeper slopes for genes N and S suggest that these genes tend to evolve faster compared to other ones. This is in line with a previous work of Dilucca et al. [24], according to which N and S are the two proteins driving the speciation process among Coronaviruses.

ORF1b has a low slope and a positive intercept in the Forsdyke plot. This points out that a number of synonymous mutations is accumulating before the phenotypic divergence. As it codes for fundamental proteins for replication, a higher conservation of the sequences is expected.

On the other hand, RdRp, that is part of gene ORF1b, has a higher slope, pointing out the faster evolution of this protein in respect to the others of the same ORF. This can be due to an selective pressure in response to antiviral drugs which mostly target this protein.

Noteworthy, all x-intercepts are very close to 0, denoting the fact that there is low accumulation of synonymous mutations in different genes, suggesting that the virus is continually evolving towards another species, due to the overall higher number of non-synonymous mutations.

### 3.7 Codon Bias and the Connectivity Patterns of SARS-CoV-2 Protein Interaction Network

As far as the network of interacting proteins in SARS-CoV-2 is concerned, we first investigated codon usage bias in relation with the degree connectivity patterns of the network. The degree distribution of the network suggests that it is scale-free (Figure 10), meaning that the network contains a large number of poorly connected proteins and a relatively small number of highly connected proteins or ‘hubs’. The corresponding genes of these hub proteins have consistently higher values of codon usage bias when this is measured by *CAI* and lower when this is measured by *ENC*, as these two codon bias indices are anti-correlated (see Table11).

**Figure 10.**
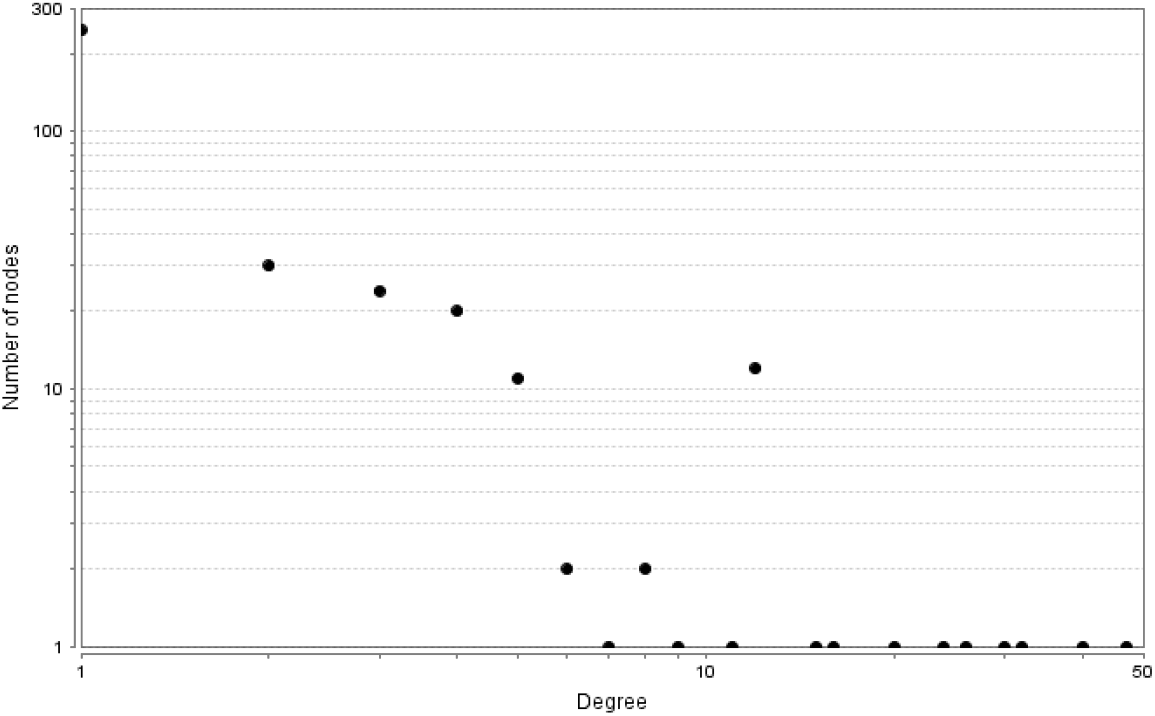
Degree distribution of proteins P(k). The degree distribution of the network follows a power law, indicating that the network is scale-free.

**Figure 11.**
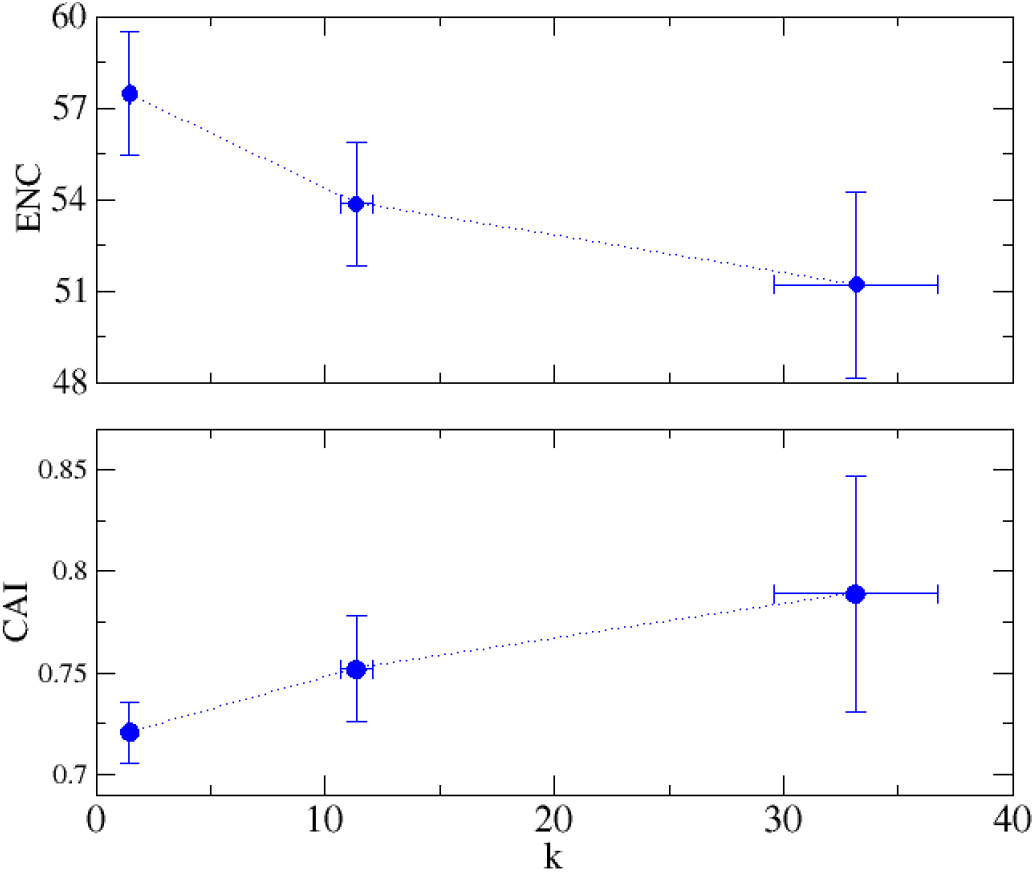
Correlation between codon bias indices of genes and the degree *k* of the corresponding proteins in the PPI. *CAI* of a gene consistently increases with the connectivity of the corresponding protein in the PPI, whereas *ENC* decreases.

Moreover, we examined codon bias in relation with the community structure of the PPI, where a community is a group of proteins that are more densely connected within each other than with the rest of the network. Table **??** shows the features of the first ten communities together with their average degree and their average of the two codon bias indices (CAI and ENC). We also calculated the internal average value 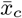 and the *Z*-scores, comparing the distribution of bias inside the community with all the proteins. Of note, all ten communities passed the Z-score test (Z*>*2). Regarding the first community, which includes only 13 proteins (3.9% of the whole network), basically overlaps with the main core of the PPI (*i*.*e*., the *k*-core with the highest possible degree). Notably, proteins belonging to this community have on average a codon bias index (as measured by *CAI* and *ENC*) that is significantly higher than the average of the rest of the network (the *Z*-score *>* 1).

## 4 Discussion

In this study we performed a comprehensive analysis of the evolutionary divergence and codon usage of SARS-CoV-2 over time, considering all genomes available in GISAID up to December 4, 2020. After filtering out incomplete genomes, we retained a total of 134,905 complete genomes, with the purpose of investigating their divergences from the first sequenced SARS-CoV-2 genome (NC 045512.2). We focused on seven SARS-CoV-2 genes/proteins that are crucial for virus structure, synthesis, transmissibility and virulence, namely the Spike, Nucleocapsid, Membrane, Envelope, *RdRp* and the two segments of *ORF* 1*ab* separately, *ORF* 1*a* and *ORF* 1*b*.

The SARS-CoV-2 genomes have a tendency to diverge constantly from the reference genome. This is in accordance with a recent study by Pachetti and colleagues (2020) where they have demonstrated that the number of SARS-CoV-2 mutations change over time [4]. The accumulation of mutations most likely affects the codon usage bias and has been investigated through RSCU, CAI, SiD and ENC.

By performing PCA in the space of RSCUs for all sequences and genes under investigation, we obtained distinct clusters: *ORF* 1*a, ORF* 1*b* and Spike exhibit very similar RSCUs and are very compact. Being also the longest genes of the genome, they represent the signature RSCU of the virus. Envelope appears to be the most spread but it should be considered that there are more than 134,000 points for that gene, that must be localized. Due to its shortness, every mutation has a high impact on the RSCU, resulting in a rather dispersed cluster.

Nucleocapsid spread instead is due to the high rate of mutations occurring in that gene.

Overall, if *E* is excluded, the genes have similar values in the major principle component (*PC*_1_), being mainly separated in *PC*_2_ that mostly accounts for 5 amino acids.

On the basis CAI and SiD time evolutions, most SARS-CoV-2 genes show a tendency to slowly decrease. This is in contrast with the initial hypothesis of adaptation of the viral Codon Usage to the host one. An explanation can be found in the already high starting values for SiD and CAI: having a close codon usage with the host would increase the efficiency of translation, for the cellular resources could be used in the most efficient way, speeding up viral replication. This can lead to the death of the host and is thought to be counterproductive for the virus, as it would stop its spreading. For this reason, an efficiency balance needs to be reached through continuous adjustments.

The ENC-plot analysis revealed that the codon usage of the viral genes under study are subject to different balances between mutational bias and natural selection. For instance, the codon preference of the genes *S, RdRp, E* and *ORF* 1*ab*, are mainly determined by natural selection, as opposed to the genes *M* and *N*, the codon usage of which is rather affected by mutational bias.

From Forsdyke plots we obtained results confirming those of our previous study [24], showing that the speciation process is predominately driven by genes *N* and *S*.

According to Rehman *et al*. (2020), the spike protein, which mediates the virus interaction with the human host cells, is more prone to mutations and particularly those occurring in the amino acids implicated in the spike-angiotensin-converting enzyme 2 (ACE2) interface [34]. The key genes, *N* and *S*, accumulate beneficial mutations that would increase the evolvability and transmissibility of SARS-CoV-2, and enable it to continuously adapt to different populations, like spilling over from bats, or other candidate natural reservoirs, to human.

RdRp, which catalyzes the transcription and replication of the coronaviral genome, binds to Nsp7 and Nsp8 to form the core RNA synthesis machinery of SARS-CoV-2 [39]. Accordingly, the genes coding for RdRp appear to be subject to natural selection, suggesting that it has an increased ability to adapt into novel hosts. Despite the fact that RdRp is considered less vulnerable to mutations [4] due to its vital role in maintaining viral genome fidelity, the mutations that occur in RdRp likely promote the virus adaptive flexibility and enhance its resistance to antiviral drugs [40].

Furthermore, we found that in the human-SARS-CoV-2 network, the most highly connected or hub proteins have consistently higher codon usage bias relatively to the less connected proteins. This observation leads to the suggestion that evolutionary pressure is exerted upon the genes encoding those proteins, most probably because of their great biological significance for the virus environmental adaptability. In other words, if these nodes are removed the entire network will eventually collapse [42].

## Footnotes

1 The density of a graph *G* with *n* nodes and *l* links is the ratio between *l* and the maximum number of possible links, namely *n*(*n−*1)*/*2, whereas, a *k*-core is a graph *G* of minimal degree *k*, meaning that each node belonging to *G* has degree greater or equal than *k*.

## Notes

### Competing Interest Statement

The authors have declared no competing interest.

